# The energy requirements of ion homeostasis determine the lifespan of starving bacteria

**DOI:** 10.1101/2021.11.22.469587

**Authors:** Severin Schink, Mark Polk, Edward Athaide, Avik Mukherjee, Constantin Ammar, Xili Liu, Seungeun Oh, Yu-Fang Chang, Markus Basan

## Abstract

The majority of microbes on earth, whether they live in the ocean, the soil or in animals, are not growing, but instead struggling to survive starvation^1–6^. Some genes and environmental conditions affecting starvation survival have been identified^7–13^, but despite almost a century of study^14–16^, we do not know which processes lead to irreversible loss of viability, which maintenance processes counteract them and how lifespan is determined from the balance of these opposing processes. Here, we used time-lapse microscopy to capture and characterize the cell death process of *E. coli* during carbon starvation for the first time. We found that a lack of nutrients results in the collapse of ion homeostasis, triggering a positive-feedback cascade of osmotic swelling and membrane permeabilization that ultimately results in lysis. Based on these findings, we hypothesized that ion transport is the major energetic requirement for starving cells and the primary determinant of the timing of lysis. We therefore developed a mathematical model that integrates ion homeostasis and ‘cannibalistic’ nutrient recycling from perished cells^16,17^ to predict lifespan changes under diverse conditions, such as changes of cell size, medium composition, and prior growth conditions. Guided by model predictions, we found that cell death during starvation could be dramatically slowed by replacing inorganic ions from the medium with a non-permeating osmoprotectant, removing the cost of ion homeostasis and preventing lysis. Our quantitative and predictive model explains how survival kinetics are determined in starvation and elucidates the mechanistic underpinnings of starvation survival.

## Results

Starving microbes undergo a gradual decrease in viability that can resemble exponential decay ^2,14,15,17–20^. While the rate and specific dynamics of the initial decay can depend on the type of starvation (Fig. S1A) and on the organism (Fig S1B), the overall phenotype of exponential decay is remarkably conserved.

To uncover the molecular mechanisms underlying this universal phenomenon, we focused on the exponential decay of viability of *Escherichia coli* K-12 in carbon starvation, where cultures lose about 99.9% of viability within 10 days (Fig. 1A). We first asked which biosynthetic processes are important for starvation survival. We measured survival dynamics in batch, while inhibiting several major cellular biosynthetic processes with different antibiotics (Fig. 1B). Strikingly, the only treatment that led to an accelerated decrease in viability and eventually extinction of the culture was the inhibition of fatty acid synthesis with cerulenin, indicating a role of the plasma membrane in starvation survival.

**Figure 1.**
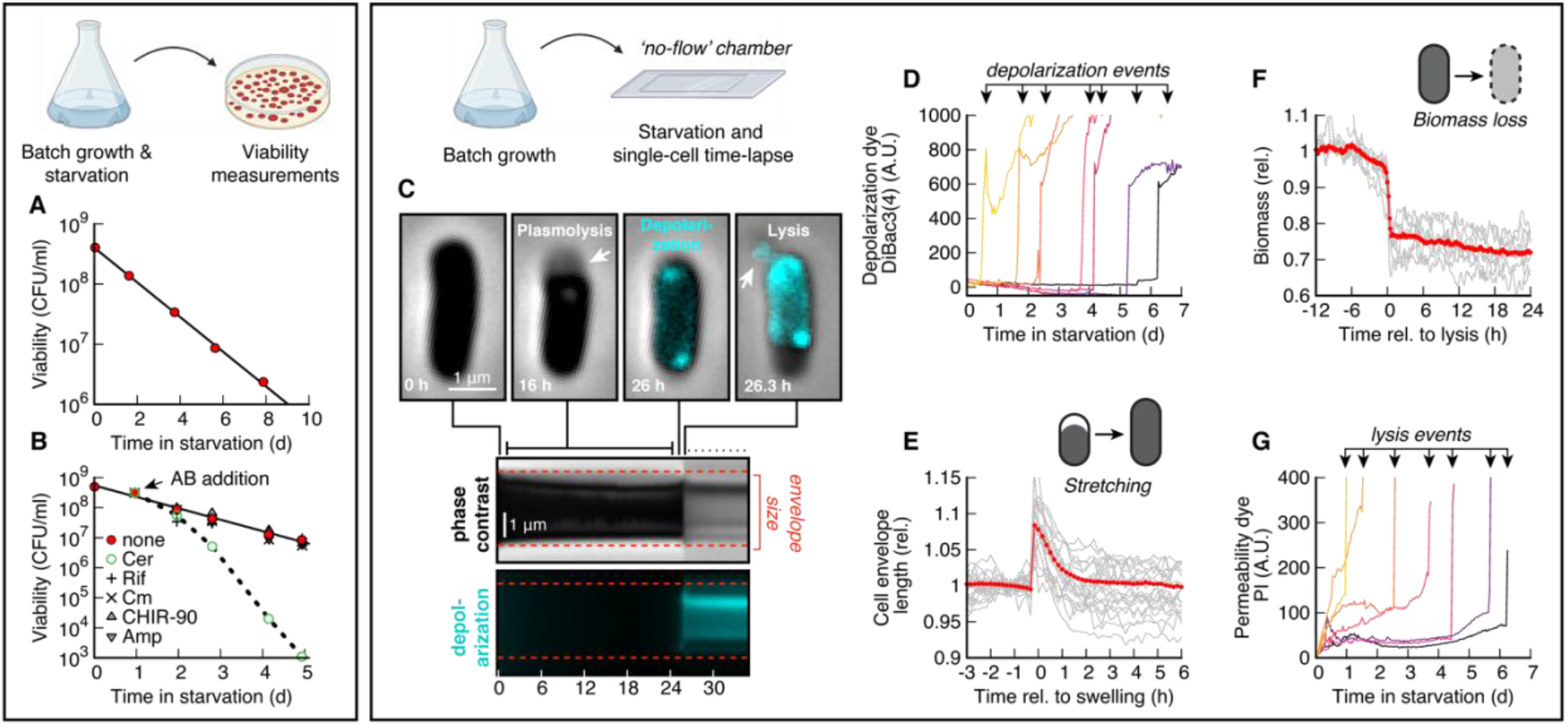
Survival dynamics of *E. coli* in carbon starvation. (**A**) Viability of *E. coli* K-12 NCM3722 in N+C- minimal medium (batch). Growth medium supplemented with 0.1% glucose; starvation induced by medium switch; viability measured in colony forming units (CFU/ml) (Methods). Line: Exponential fit. Data points are an average of n=3 replicates. (**B**) Viability of control (red) compared to cerulenin (Cer, fatty acid synthesis), rifaximin (Rif, transcription), chloramphenicol (Cm, translation), CHIR-90 (lipid-A synthesis) and ampicillin (Amp, cell wall synthesis). Antibiotics were added daily, starting at 24 h at 10x MIC. Data points are an average of n=3 replicates. (**C**) Live-cell imaging of *E. coli* in a ‘no-flow’ chamber (Methods). Plasmolysis, the retraction of cytoplasmic membrane from cell envelope, lasted from the first hours of starvation until shortly before death (see Kymograph). During plasmolysis, the cytoplasm spontaneously depolarizes (cyan DiBac3(4) signal turns on at 26 h), which coincides with swelling (Fig. S2E&F). Finally, the bacterium lyses (‘arrow’ at 26.3 h), which is typically delayed from swelling (Fig. S2G-H). Total # of bacteria: 14.498. (**D**) Single cell traces of seven representative bacteria dying on different days show a sudden spike signal, indicating rapid loss of ion homeostasis. (**E**) Cell envelope length of single cells, aligned to time-point of swelling (‘grey’; mean in ‘red). # of bacteria: 858. (**F**) Single-cell biomass, measured with quantitative phase microscopy (QPM; Methods) aligned to time-point of lysis. Loss of biomass: 28±7 % (Fig. S2G). # of bacteria: 34. (**G**) Single-cell traces of fluorescence signal of propidium iodide (PI) of the seven cells of panel C. Permeability is measured as the slope of the staining dynamics. See Fig. S4 for combined DiBac3(4) and PI traces of individual bacteria.

To determine the causes of loss of viability in starvation, we imaged individual bacteria during starvation with live-cell, time-lapse microscopy, using a membrane potential dye as a readout for ion homeostasis; see Methods. We found that within a few hours after nutrient depletion, the cytoplasm of *E. coli* contracted and detached from the cell wall at the poles (Fig. 1C, image at 16 h; Fig. S2A&B - quantification and statistics, Fig. S2C&D - SEM & TEM), a process called plasmolysis that is typical for bacteria in starvation^18,21^. By continuing to track starving bacteria for seven days, we captured the death process in thousands of bacteria: spontaneous depolarization of the cytoplasm was rapidly followed by swelling and lysis (Fig. 1C, 26 h & 26.3 h; Fig. 1D). Swelling of the cytoplasm led to stretching of the cell envelope, indicated by an increase in cell length (Fig. 1E, Fig. S2E&F), and lysis led to a sudden loss of biomass (Fig. 1F, Fig. S2G&H). Analyzing the succession of events, we found that the collapse of membrane potential coincided with cytoplasmic expansion (−0.05±0.50 h, Fig. S2F) and preceded lysis (2.0±1.4 h, Fig. S2H). This led us to hypothesize that the uncontrolled expansion is caused by a catastrophic failure of ion homeostasis, resulting in a build-up of osmotic pressure and eventually lysis. We next stained bacteria with propidium iodide (PI), a non-toxic stain (Fig. S3A) that intercalates irreversibly with DNA (Fig. S3B&C), which allows us to use it as a metric for cell permeability (see Fig. S3D, E). We found that PI diffuses through the membrane at a slow rate during starvation, indicating that the membrane is slightly permeable in viable bacteria, but shows a sharp increase when bacteria lyse (Fig. 1G).

To investigate the morphology of the cell envelope, and in particular the inner membrane, we also performed transmission electron micrography (TEM) during starvation survival. Electron microscopy revealed dramatic changes of membrane morphology. While early in starvation there was excess cytoplasmic membrane, after 48 h in starvation the excess material had vanished and the shape of the membrane at the pole was nearly half-spherical, indicating tension on the membrane (Fig. S2C).

Based on these observations, we hypothesized that maintaining ion homeostasis and membrane integrity are the two most important processes that prevent lysis of starving bacteria. The role of membrane integrity is straightforward to understand, but the role of ion homeostasis is less obvious. However, all cells must actively transport inorganic ions out of their cytoplasm to reduce their internal solute concentration. If this process fails, the excess concentrations of metabolites, macromolecules and counterions inside the cell leads to the influx of water, swelling and lysis^22,23^. Therefore, maintaining osmolarity to prevent swelling is a continuous process that consumes energy that is in short supply during starvation. If our hypothesis is correct, this would suggest that maintenance of ion homeostasis is the largest indispensable energy cost of starving cells.

To test whether these ideas are sufficient to explain the key experimental observations on starving bacterial populations, we devised a simple mathematical model (Fig. 2A). One important goal for the model was to explain how the exponential decay of viability of Fig. 1A emerges from the seemingly random time to death of single cells, as observed in Fig. 1D. The model assumes that ions from the medium (red, coarse-grained into a single species) diffuse into the cytoplasm and are exported using active transport, which consumes energy provided by nutrients (green); see Eq. (1). Nutrients released from dying cells are cannibalized and provide an energy source to surviving cells^16,17^; see Eq. (2). Based on experimental findings^24–26^, we also assumed that membrane permeability is increased by membrane tension. Biological fluctuations in permeability and pumping are integrated in a normal-distributed noise term.

**Figure 2.**
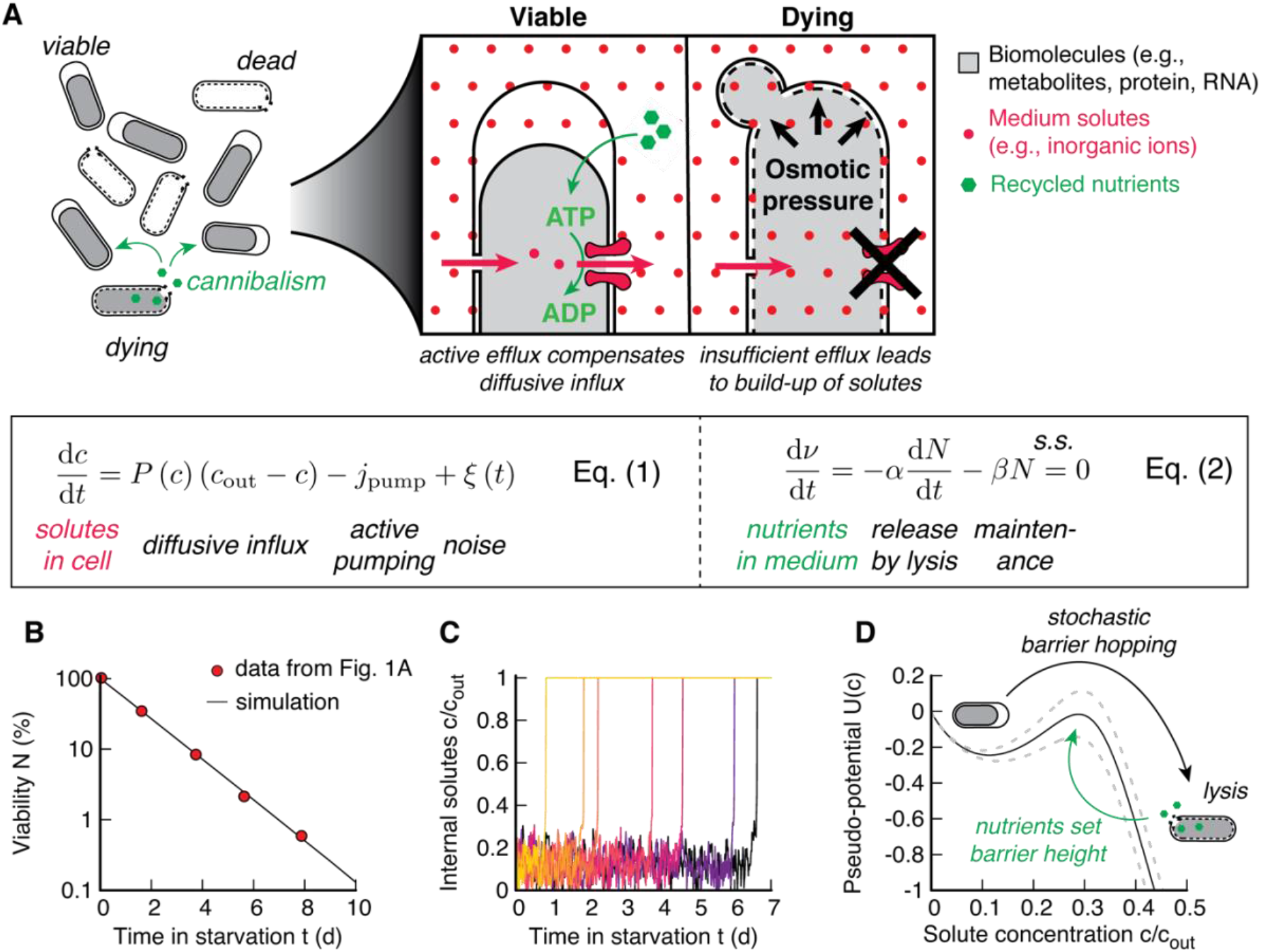
Ion homeostasis maintenance model. (**A**) Nutrients released by lysis are cannibalized by viable bacteria (left) and are used to pump out solutes (e.g., ions) from the cytoplasm to the medium (right). If sufficient nutrients are present to support pumping, then the cytoplasm contracts (plasmolysis) until the concentration of solutes from biomass balances *c_out_*. Insufficient pumping leads to collapse of ion gradients, osmotic water inflow and swelling, resulting in lysis. The dynamics of internal solute concentration, *c_in_*, (Eq. 1) are given by diffusive influx (Fick’s law; permeability *P*, ion gradient Δ*c* = *c_out_* – *c_in_*), active pumping (*j_pump_*, which is a part of the total maintenance *β*), and normal-distributed noise *ξ*(*t*). Stretching of the membrane by increased turgor pressure (i.e., decreased Δ*c*) increases *P*(Δ*c*). The dynamics of nutrients in the medium *v* is given by release of nutrients (recycling yield *α*), and maintenance *β*. N is the number of viable bacteria in the culture. In steady state ‘s.s.’, *v* = *const*. See SI for details on the model. (**B**) Simulated loss of viability of 10^5^ bacteria (black line) compared to experimental data from Fig. 1A. (**C**) Dynamics of single-cell solute concentration for bacteria dying at different time-points shows random variations around a small solute concentration, and a spontaneous loss of homeostasis, analog to Fig. 1D. (**D**) The model can be mapped onto a potential *U*(*c*), ‘black line’, with *dc*/*dt* = –*U*′(*c*) + *ξ*(*t*). Height of the barrier self-adjusts. Dashed lines show the effect of a 10% change in death rate on the potential barrier.

Simulations of this ‘ion homeostasis’ model closely matched experimental observations. Indeed, the model dynamics yielded an exponential decay in viability (Fig. 2B) and a spontaneous loss of ion homeostasis and death (Fig. 2C). This is because in the model, the spontaneous death process can be understood as a stochastic escape from a potential well (Fig. 2D), similar to the classic Kramer’s escape model in physics^27^. In the potential well, starving cells are in plasmolysis, minimizing tension on the membrane. But unlike in the classic Kramer’s escape model, the potential barrier itself is created by nutrients released from dying cells, i.e., cells that have crossed the potential barrier. This leads to a steady state where most bacteria are supplied with just enough nutrients to remain in the potential well, but the potential barrier remains small enough to let cells to escape the well at a low rate and provide nutrients to support surviving cells. Therefore, because the barrier height itself is set by the escape rate over the barrier, even small fluctuations in the underlying biological variables are sufficient to result in a highly stochastic exponential decay in viability; see SI. This closely resembles our experimental observations, where the vast majority of starving cells remain in plasmolysis (low internal ion concentration), but a subset stochastically undergoes expansion and lysis (ion influx and crossing the potential barrier).

To experimentally test this model and the ion homeostasis mechanism it describes, we derived a set of quantitative predictions from the model. For bacteria that are in the potential well, the internal ion concentration *c*_in_ is in steady-state and ion efflux from active transport, *j*_out_, is balanced by passive diffusive influx, *j*_in_. Fick’s law of diffusion states that the diffusive influx *j*_in_ is proportional to the concentration gradient of ions across the membrane, *j*_in_ = *P*Δ*c*, where *P* is the permeability of the membrane and Δ*c* = *c_out_* – *c*_in_ is the ion gradient. Hence, the energy expenditure of bacteria for pumping ions, denoted by *β*, is given by, *β* = *κj*_out_ = *κP*Δ*c*, where *κ* is the pumping cost. On the other hand, energy for this active transport process comes from nutrients released by dying cells. If the death of a cell yields *α* nutrients that are divided up equally among all *N* viable bacteria, and 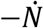 are dying per unit time, then the rate at which a bacterium is supplied with nutrients is 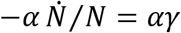, where *γ* is the death rate. In nutrient-recycling steady-state, *v* = 0, nutrient supply and nutrient demand balance, so *αγ* = *β* and we obtain a quantitative relation for the death rate as a function of permeability, ion gradient and the recycling yield

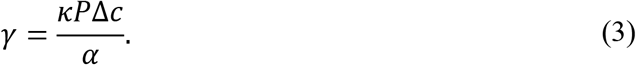

In addition, in this simplified one-ion model, we assume that viable bacteria keep *c*_in_ low and independent of *c*_out_ and *P*, based on the observation that bacteria keep concentrations of unwanted ions low in a wide range of media, and balance osmotic pressure primarily with glutamate and potassium^28^.

Eq. (3) makes predictions about how lifespan should scale with cell size. Permeability *P* is proportional to the cell surface area *S*, because diffusion happens across the cell membrane, while the recycling yield *α* is proportional to cell volume *V*, because bigger cells release more nutrients^20^. Thus, according to Eq. (3), death rate should scale with the surface-to-volume ratio of bacteria, *γ* ∝ *S/V*, if all other factors are kept constant. There are plausible alternative hypotheses for the reasons for cell death that would result in markedly different scaling. For example, if death was caused by a single, rare catastrophic event, such as damage of the cell membrane or accumulation of a toxic protein aggregate, death would be more likely for larger cells. Thus, we would expect scaling of death rate to be proportional to membrane material, *γ* ∝ *S*, or cytoplasmic material, *γ* ∝ *V*. On the other hand, if death is caused by a gradual deterioration of either the cell’s membrane or its cytoplasmic material, a bigger cell would survive longer and death rate would decrease proportional either to the inverse of membrane material, *γ* ∝ *S*^−1^, or to the inverse of the total cytoplasmic material, *γ* ∝ *V*^−1^.

To test whether these alternatives are better explanations of cell death during survival than our ion homeostasis model, we varied cell length of *E. coli* using a titratable expression construct of the cell division protein FtsZ^29^ (Fig. S6). Remarkably, we observed an almost constant death rate, despite a 2.5-fold variation in average length (Fig. 3A) that is clearly incompatible with the scaling expected from the alternative models (black lines). Moreover, FtsZ-deprived cells also typically contain several copies of chromosomal DNA^30^, which suggests that DNA damage is unlikely to be the death mechanism. Instead, the slight decrease of death rate in longer cells closely matches the estimated surface-to-volume ratio of rod-shaped bacteria, precisely as predicted by Eq. (3) (Fig. 3A, red line).

**Figure 3.**
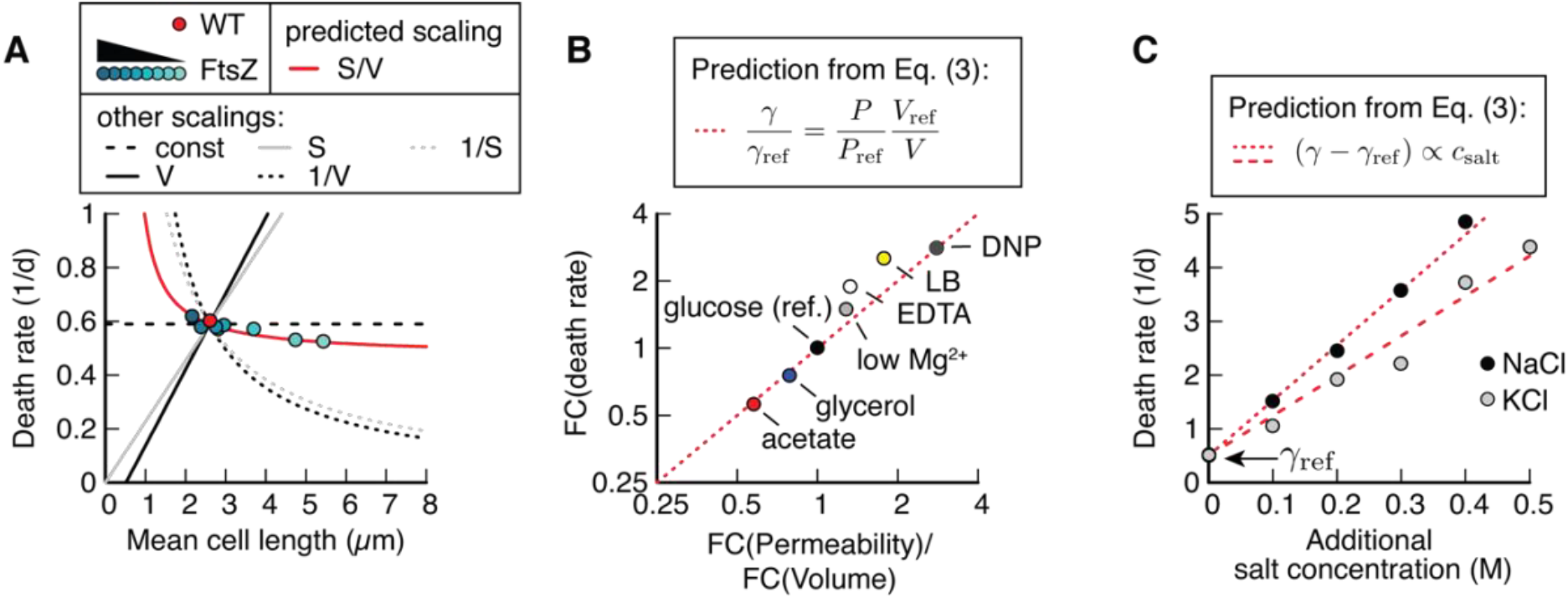
Cost of ion homeostasis determines death rate. (**A**) Death rate of wild-type and mutant ptac-ftsZ^27^, induced with varying levels of IPTG, resulting in a change in cell length (Fig. S6). Change of death rate compared to the predicted S/V scaling, and proportional or inverse scaling with either S or V. (**B**) Fold-change (FC) of death rate, measured in batch cultures plotted versus foldchange of mean permeability (Methods) times the inverse fold change of volume, both measured in time-lapse microscopy. Theoretical prediction: Unity line in dashed red. Reference condition: growth in minimal medium with glucose (black). Red, blue: supplemented with different carbon substrates (acetate or glycerol) during growth. Light-grey: Mg^2+^ is changed during starvation from 0.4 mM to 0.1 mM. White: 1 mM EDTA added in starvation. Yellow: Growth on LB. Dark-grey: 1 mM DNP (di-nitrophenol) added in starvation. Death rates are measured as average of n=3. Permeability and volume changes are the average of 2.000 to 20.000 individual bacteria. (**C**) Death rate of cultures supplemented with either KCl or NaCl during starvation. Fit: *R*^2^ = 0.99 (NaCl), *R*^2^ = 0.96 (KCl).

To further investigate the ion homeostasis model’s predictions, we wondered if differences in death rates that are observed when bacteria are grown in different conditions could be explained by Eq. (3). Different growth conditions prior to starvation are known to affect maintenance rate for unknown reasons^20^. These changes in death rate are not accounted for by a change in *S/V* because *S/V* decreases for faster growth (because cells become bigger), while death rate increases. Based on Eq. (3), we suspected that these changes in death rate might instead be caused by changes in bacterial permeability. To test this, we cultured *E. coli* in different growth media, washed them and resuspended them in carbon free medium. From Eq. (3), the fold change (FC) of death rate should be reflected in the fold changes of permeability and volume, FC(*γ*) = FC(*P*) FC^−1^(*V*). Indeed, quantifying average permeability for these starvation conditions, using the slope of PI staining (Fig. 1F) between 0 and 10 h, showed that the change of death rate in slow-growth carbons and rich media could be explained solely by the effect on permeability and volume (Fig. 3B). To further test the causality of the effect of permeability on death rate, we permeabilized *E. coli* with low Mg^2+^, EDTA or DNP. These higher permeabilities were reflected in faster death rates (Fig. 3B). Death rates for all experimental conditions closely followed the model prediction of Eq. (3) (Fig. 3B, identity line in red).

Finally, the ion homeostasis maintenance model makes the prediction that death rate should depend on the medium composition. According to Eq. (3) increasing the external ion concentration *c*_out_ should linearly increase death rate because an increase in ion gradients leads to an increase in diffusive influx, which needs to be balanced with a higher pumping rate to keep the internal ion concentration low. Indeed, as predicted, supplementing the medium with either NaCl or KCl led to a striking linear increase in death rate, up to a 10-fold increase at 0.5 M for both salts (Fig. 3C).

For a final test, we asked if it would be possible to reduce death rates by lowering salt concentrations in the medium. Consistent with our earlier observations, decreasing the concentration of our phosphate-buffered minimal medium (N+C-) decreased death rate, but only by about a third, before increasing again at low osmolarity (Fig. S7). This can easily be understood by considering that a minimum level of external osmotic pressure is crucial to balance the internal osmotic pressure from metabolites, biomolecules and cellular counterions, in order to prevent cell swelling.

Guided by our model, we reasoned that it should be possible to overcome this limitation of cell survival in low-salt medium by balancing cellular osmotic pressure with external osmotic pressure from an impermeable osmolyte. The low-salt medium should minimize the energy cost required to pump out ions, while the cellular osmotic pressure would be balanced by an external osmolyte to prevent lysis. We therefore devised a new ‘osmo-balanced’ medium that consists of 0.2 M MOPS, an inert and impermeable buffer molecule, and low quantities of essential ions; see Methods. We grew *E. coli* in our regular medium to ensure an identical cell state before starvation, then washed and resuspended the bacteria in the ‘osmo-balanced’ medium. Remarkably, this medium led to a 4-fold decrease in death rate compared to our regular culture conditions (Fig. 4A). This improvement of survival was not unique to *E. coli;* other bacterial species showed similarly improved starvation survival in our osmo-balanced medium (Fig. S1C&D). Together, the findings in Fig. 4A and Fig. 3C demonstrate that starvation survival can be controlled across a more than 40-fold range of death rate, simply by changing medium salt concentrations.

**Figure 4.**
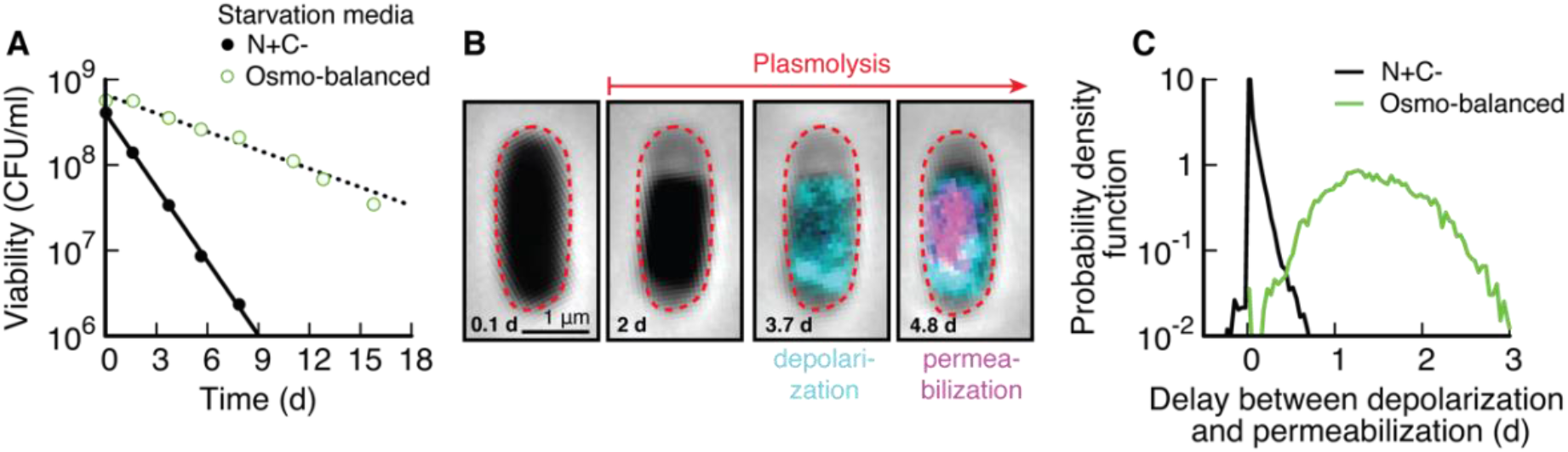
Rescued survival dynamics in ‘osmo-balanced’ medium. (**A**) Comparison of viability (batch) of cultures grown in N+C+ and starved in either N+C- (black, data from Fig. 1A.) or ‘osmo-balanced’ medium (green). Death rate decreased 4.0-fold. n=3. (**B**) Time-lapse images of *E. coli* in ‘osmo-balanced’ medium. After depolarization, the cytoplasm does not swell and the bacterium does not lyse. (**C**) Delay distribution between depolarization and permeabilization. In N+C- medium (‘black’) both events are in rapid succession, while in ‘osmo-balanced’ medium (‘green’) the events are disconnected by 1.5±0.5 days. In addition, in the ‘osmo-balanced’ medium both depolarization and permeabilization are gradual (Fig. S8), compared to the rapid rate of both processes in N+C- (Fig. S4). N+C-: 14.498 Osmo-balanced: 11.023 bacteria.

To confirm that our ‘osmo-balanced’ medium prevented cytoplasmic swelling, we performed live-imaging of *E. coli*. Several days into starvation, bacteria began to depolarize, presumably due to lack of energy. However, depolarized bacteria remained in plasmolysis and did not swell and lyse (Fig. 4B). Most bacteria eventually permeabilized, but only several days after the loss of membrane potential, as opposed to the two events coinciding, as occurred in our regular culture medium (Fig. 4C). Depolarization and permeabilization happened gradually (Fig. S8) rather than abruptly as in regular minimal medium (Fig. S4). Thus, the osmo-balanced medium disrupted the normal death process and decoupled depolarization from osmotic swelling and lysis. Unsurprisingly, bacteria did not survive indefinitely in the osmo-balanced medium, which indicates that while ion homeostasis is the major energy-consuming process in starvation, secondary processes also play a role in survival, for example membrane turnover (see Fig. 1B).

Our results show that maintenance of ion homeostasis is the major energetic cost limiting bacterial survival in starvation. Moreover, failure of ion homeostasis, triggered by a lack of energy, is the cause of death of starving bacteria. The ion homeostasis maintenance model introduced here explains how the stochastic exponential survival dynamics of single bacteria during starvation emerge from the collective phenomena of maintenance and nutrient exchange. The model enables quantitative predictions of how survival rates are affected by diverse environmental conditions and bacterial adaptations. Based on our findings, we propose that targeting ion homeostasis could be a promising avenue for developing novel antibiotics against non-growing and dormant infections, for which currently available antibiotics are often ineffective. Finally, because most microbes on earth spend most of their time in starvation conditions, we expect the principles underlying starvation survival elucidated in this work, to be of broad relevance for understanding the evolutionary strategies that contribute to the striking diversity of phenotypes of microbes and microbial communities in different natural ecological niches.

## Acknowledgements

This project was supported by MIRA grant (5R35GM137895) and an HMS Junior Faculty Armenise grant to MB. SJS was supported by EMBO via a Long-term fellowship (ALTF 782-2017) and HFSP via a long-term fellowship (LT000597/2018). MP was supported by the Harvard College Research Program (HCRP). EA was supported by HCRP and the Program for Research in Science and Engineering (PRISE) of Harvard. XL and SO were supported by the NIH grant (R56AG073341). Microscopy was performed at the Nikon Imaging Center at Harvard Medical School. Electron microscopy was performed at the Center for Nanoscale Systems at Harvard University. We thank Suckjoon Jun for sharing strain VIP205, Marc Kirschner for helping us with QPM microscopy and Rebecca Ward for many helpful suggestions.

## Data availability

All analyzed data are available in the supporting information. Raw microscopy data can be shared upon request.

## Contributions

SJS and MB conceived this study, designed experiments and theory. SJS performed the majority of experiments with contributions from MP in starvation experiments and microscopy, and AM in electron microscopy and QPM imaging experiments. SJS, EA and MB performed modelling. SO contributed to the QPM instrumentation and SO and XL designed the QPM image processing pipeline. YFC constructed strains. CA contributed data analysis and ideas on membrane integrity. SJS and MB wrote the paper with input from MP, AM, EA, CA, XL & SO.

## Methods

### Strains

All strains used in this study are derived from wild type *E. coli* K-12 strain NCM3722^31^. To titrated FtsZ, the construct ptac::FtsZ of strain VIP205^29^ was transferred to NCM3722 via P1 transduction. Bacteria used in Fig. S1 are environmental isolates kindly provided by the lab of Roberto Kolter.

### Culture medium

In this work we used two distinct culture media. Our standard medium was N-C- minimal medium^32^, containing K2SO4 (1 g), K2HPO4·3H2O (17.7 g), KH2PO4 (4.7 g), MgSO4·7H2O (0.1 g) and NaCl (2.5 g) per liter. The medium was supplemented with 20 mM NH4Cl, as nitrogen source, and 0.1% glucose as the sole carbon source, unless indicated otherwise. For carbon or nitrogen starvation, N-C- minimal medium was prepared either without carbon (N+C-) or nitrogen supplement (N-C+), respectively. The second medium is our ‘osmo-balanced’ medium, which contains 0.2 M MOPS (3-(N-morpholino)propanesulfonic acid), titrated to pH 7 with KOH, and 1 mM MgCl_2_, 0.1 mM CaCl_2_, 0.16 mM K_2_SO_4_, 0.5 mM K_2_HPO_4_, 22mM NH_4_Cl and lacks a carbon source. Cultures starved in the ‘osmo-balanced’ medium were previously grown in N-C- medium supplemented with NH_4_Cl and glucose.

### Growth protocol

Growth was performed in batch cultures in glass test tubes (Fisher Scientific) with disposable, polypropylene Kim-Kap closures (Kimble Chase) in a water bath shaker kept at 37°C. Single colonies were picked from an LB agar plate, grown in LB until saturation and used to inoculate overnight cultures in N-C- minimal media supplemented with NH4Cl and glucose. Dilutions were chosen such that cultures did not saturate overnight to ensure continuous exponential growth. The next day, cultures were diluted into fresh minimal medium and grown for at least 5 doublings. Starvation was induced by centrifugation (5000 RCF for 3 min), removal of supernatant and resuspension in fresh, carbon-free N-C- minimal medium or ‘osmo-balanced’ medium. This protocol was chosen to avoid bacteria using waste products such as acetate after glucose was depleted^33^. For nutrient sources without acetate excretion (e.g. glycerol), this washing step does not alter the survival kinetics^17^. To achieve ‘stationary phase’ adaptation we used the effect of excreted waste products and let *E. coli* grow on glucose until depletion, followed by washing 12 h to 24 h later.

### Viability counting

Samples were diluted in fresh N-C- minimal medium without carbon substrate to an estimated target density of 4000 CFU/ml using a multi-channel pipette and a 96 well plate. 100 μl of the diluted culture were spread on LB Agar using Rattler Plating Beads (Zymo Research) and incubated for 24 h. LB Agar was supplemented with 25 μg/ml of 2,3,5-triphenyltetrazolium chloride to stain colonies red and increase contrast for automated colony counting using an automated CellProfiler pipeline^34^. Images were taken on a Rebel T3i (Canon) mounted on top of a Lightpad A920 (Artograph).

### Live-cell Microscopy

To image individual bacteria for seven days, we designed a ‘no-flow’ chamber that minimizes nutrient contamination and evaporation. The chamber consists of a sandwich of a cover slip (22×50 mm, No 1.5, VWR) and a microscope slide (3”x1”x1mm, VistaVision, VWR), with a thin layer of PDMS in-between. At the top and bottom, the culture is in contact with inert glass that prevents evaporation or leakage of nutrients from the plastic. Two holes are laser-cut into the microscope slide (Universal Laser Systems PLS 6.150D) through which the chamber will be filled. The PDMS layer (Sylgard 184, 1:10 ratio, Dow Corning) is spin coated on a silicon wafer for 30 s at 500 rpm (Laurell, WS-650MZ-23NPPB), cured at 85°C for 1 h, peeled off and gently draped over the microscope slide. Then, the incubation chamber is cut out of the PDMS with a laser cutter and unwanted PDMS pieces are removed from the microscope slide with tweezers. The resulting microscope slide with PDMS chamber is plasma treated 50 W, 10 s, O2, duty ratio 255 (Tergeo Plasma Cleaner, Pie Scientific) and the cover slide is added on top. The finished chamber is cured overnight at 85°C.

Cultures taken from exponential growth at OD 0.5 are starved according to the above protocol and stained with 5 μg/ml DiBac3(4) and 1.25 μg/ml Propidium Iodide (Thermo Fisher Scientific) 10 minutes after entry to starvation. The ‘no-flow’ chamber was then filled with the stained, starved culture through the laser-cut holes with a starved culture at OD 0.5, identical to batch experiments. Holes are then sealed (AlumaSeal, Sigma-Aldrich) and the chamber is centrifuged, cover slide side down at 1200 RCF for 2 min, to adhere bacteria to the cover glass. The majority of bacteria stably adhered to the glass surface for seven days without coating. Apart from the filling and centrifugation period, which were at room temperature, the culture and chamber were kept at 37°C throughout the process.

Phase contrast and fluorescence microscopy was performed on a Nikon Ti-2 equipped with a 100× phase contrast immersion oil objective, Hamamatsu Flash 4.0 sCMOS camera, Lumencor Spectra-X light engine and OkoLab cage microscope incubator set to 37°C. To image the time-lapse, Nikon Elements was used to control illumination times (phase: 200ms, green 10 ms, red 10 ms.) and illumination strength (phase: 100%, green 10%, red 10%). A combination of perfect focus, a z-stack (4 steps of 0.4 μm) and a software autofocus (post-acquisition) were used to obtain continuous images of bacteria in focus.

Single cell dry mass was measured using quantitative phase microscopy (QPM) on a Nikon Eclipse Ti with a Nikon, CFI Plan Apo Lambda 100X objective with 1.5 N.A. an 2x magnifier lens (Thorlabs, EX2C extender) and a commercial phase sensor (Phasics, SID4bio). In the trans-illumination light path, a colored-glass bandpass filter (FGB39, Thorlabs) was mounted to select the blue light in order to increase the spatial resolution. This phase sensor enables single-shot acquisition of quantitative phase through quadriwave lateral shearing interferometry (QWLSI)^35^ by using a 2-dimensional diffraction grating (modified Hartmann mask) positioned at a short distance from a CCD sensor. The raw image of quantitative phase, measured as optical path difference (OPD), is induced by both the sample and the optical system itself. We used the ceQPM method as reported previously^36^ to obtain the OPD of the optical system and fit the background fluctuation that are then subtracted from the raw image to obtain the OPD of the cell. The dry mass *m* is computed from the cell’s OPD through the relation *m* = 1/*α* ° ∮ OPD(*x*, *y*)d*x*d*y*, where *α* = d*n*/d*c* is the average refractive index increment of cellular materials and the integral is over the cell area S. We used α = 0.18 ml/g to convert the OPD to dry mass, which is the typical value for most cellular materials^37^.

Time-lapse was analyzed in MATLAB. First, regions of interests (ROIs) with an object in phase contrast resembling a bacterium were identified. Next, for every time-point, the correct focal plane was identified, the fluorescence background was subtracted, and the ROIs were cut out and saved separately. Finally, ROIs were analyzed and the total fluorescence (by summing up the value of all pixels), as well as length, width and area of the individual bacteria were recorded. For QPM analysis, the segmented bacterium was dilated and the total phase-shift was integrated to yield the total biomass. This analysis yielded 14,500 single-cell time-traces of bacteria starving in N-C- medium (Fig. 1) and 11,000 for the ‘osmo-balanced’ medium (Fig. 4).

### Scanning electron microscopy protocol

Culture was harvested by filtering 2 ml of exponentially growing or starving bacterial culture at 0.5 OD through a 0.22-micron filter (Durapore 0.22 μM PVDF membrane, Millipore GVWP02500) held on a vacuum micro-filtration device (25mm glass filter holder, Millipore XX1012502). Immediately after this step, the filter was submerged in Karnovsky fixative (Electron microscopy sciences 15732-10). Cells were dislodged from the filter by repeated but gentle pipetting with 1 ml pipette and kept in fixative solution for 1 hour at room temperature. After one hour, 500 ul of fixative containing cells were added on 12 mm glass coverslip kept on a 35mm petri plate (glass coverslips were precoated with Ploy-L-Lysine) and centrifuged in a swing bucket centrifuge (Eppendorf) for 10 min at 5000 g. After centrifugation, the glass coverslips with adherent cells were placed in histology tissue fixation and processing cassettes (electron microscopy sciences) and the cassettes were sequentially dipped in graded series of ethanol (30%, 50%, 75%, 80%, 90%, 95% and 100%). Grades of ethanol were prepared using anhydrous ethanol, 200 proof obtained from electron microscopy sciences (15058). Cells were kept in each grade of ethanol for 10 min, and the 100% ethanol step was repeated three times with fresh ethanol to ensure perfect dehydration. After this step cells were kept submerged in 100% ethanol overnight (at 4°C) and taken to Harvard center for nanoscale systems (CNS) next morning for further processing. Prior to microscopy, cells were critical point dried in a Tousimis critical point dryer, mounted on SEM sample holder and coated with heavy metal (gold/palladium) using sputter coater (Hummer) and imaged in JEOL 5600 LV scanning electron microscope. Processing steps at CNS and imaging was aided by the CNS stuff and facility in charge.

### Transmission electron microscopy protocol

For transmission electron microscopy, bacterial cell samples were collected similarly as scanning electron microscopy method. Briefly, 2 ml of bacterial culture at 0.5 OD was taken and cells were harvested by filtration. Filter containing harvested cells were immediately dipped in Karnovsky fixative, and cells were dislodged from filter by gentle pipetting. Cells in fixative were taken to CNS for further processing. At CNS, cells in fixative were centrifuged to obtain a cell pellet, and the pellet was embedded in resin and thin sections were prepared. Cut sections were mounted on EM grid and imaged using HT7800 Hitachi TEM system.

### Quantification of permeability

Live-cell microscopy with propidium iodide staining, as described see above, was used to measure single-cell permeability. Fluorescence signal was integrated over the entire area of each cell, as identified by thresholding in phase contrast, to calculate the total PI staining per cell of each cell per time-point. PI staining rate of each cell was fitted as a direct proportionality in the period from 0 to 10 h to calculate the total permeability across the membrane. To calculate the change of permeability compared to the reference condition, we used a ‘no-flow’ chamber with 6 parallel channels, that allows us to measure permeability of five conditions and the reference condition in the same experiment.

### Quantification of single-cell parameters

To quantify plasmolysis, the threshold for segmentation of the bacterium was chosen such that it contains only the cytoplasm (dark area). To quantify the length of the cell envelope, the threshold was increased, such that the cell envelope (light grey are) was included in the segmentation. Both thresholding techniques are semi-quantitative, as the exact boundary of the outer and inner membrane cannot be resolved with phase microscopy, but they circumvent the problems generated by potentially toxic membrane stains.

The Fluorescence signal from DiBac3(4) and propidium iodide was integrated over the entire segmented bacterium and normalized to the cell area to correct for differences in cell size. The time-point of depolarization and permeabilization was determined by the first time the signal reached a threshold. For depolarization, the threshold was chosen to be the local minimum of the bimodal histogram of DiBac3(4) signals of all cells throughout seven days of starvation. For propidium iodide, the threshold was chosen manually. The threshold is indicated in the example dynamics of individual cells in Fig. S4 and S8.

To align single-cell stretching and biomass loss, we searched in the vicinity of the time-point of depolarization for the largest increase in length and the largest decrease in biomass, respectively. The time delay between the time-point of depolarization and the time-points of stretching and biomass loss is reported in Fig. S2.

### Cell size

Cell volume was computed as *V* = *π*(*w*/2)^2^(*l* – *w*) + 4/3*π*(*w*/2)^3^ where *w* and *l* are width and length of each cell, whose shape was considered as a cylinder (with radius *w*/2 and height equal to (*l* – *w*)) with two semi-spheres at the ends (with radius *w*/2). Surface area was calculated analogously as S = 4*π*(*w*/2)^2^ + *w*/2 (*l* – *w*). Because cell width only changes marginally in the FtsZ titration, we assumed it to be constant and took the formulas of *S* and *V* at wild-type width to plot the length dependence of surface and volume in Fig. 3.

