## Supplementary Material for "The energy requirements of ion homeostasis determine the lifespan of starving bacteria"

#### Supplementary Figures

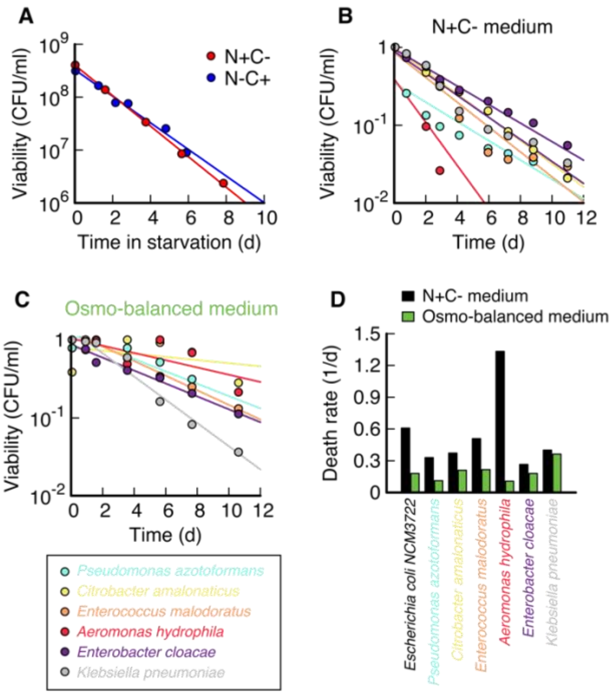


**Figure S 1 Starvation of other substrates and of other bacterial species.** (**A**) Nitrogen starvation (blue) and carbon starvation (red) of E. coli K-12 NCM3722 in N-C+ and N+C- media, respectively (Methods). (**B**) Six gram-positive and gram-negative bacteria starved in N+C- medium, with previous growth on N+C+ with 0.1% glucose. (**C**) Starvation of the same bacteria as panel B, but in 'osmo-balanced' medium, identical to Fig. 4. (**D**) Comparison of death rates in N+C- and 'osmo-balanced' media. In all bacteria death rate in the 'osmo-balanced' medium is slower than in N+C-, with the effect size being different for individual bacterial species.

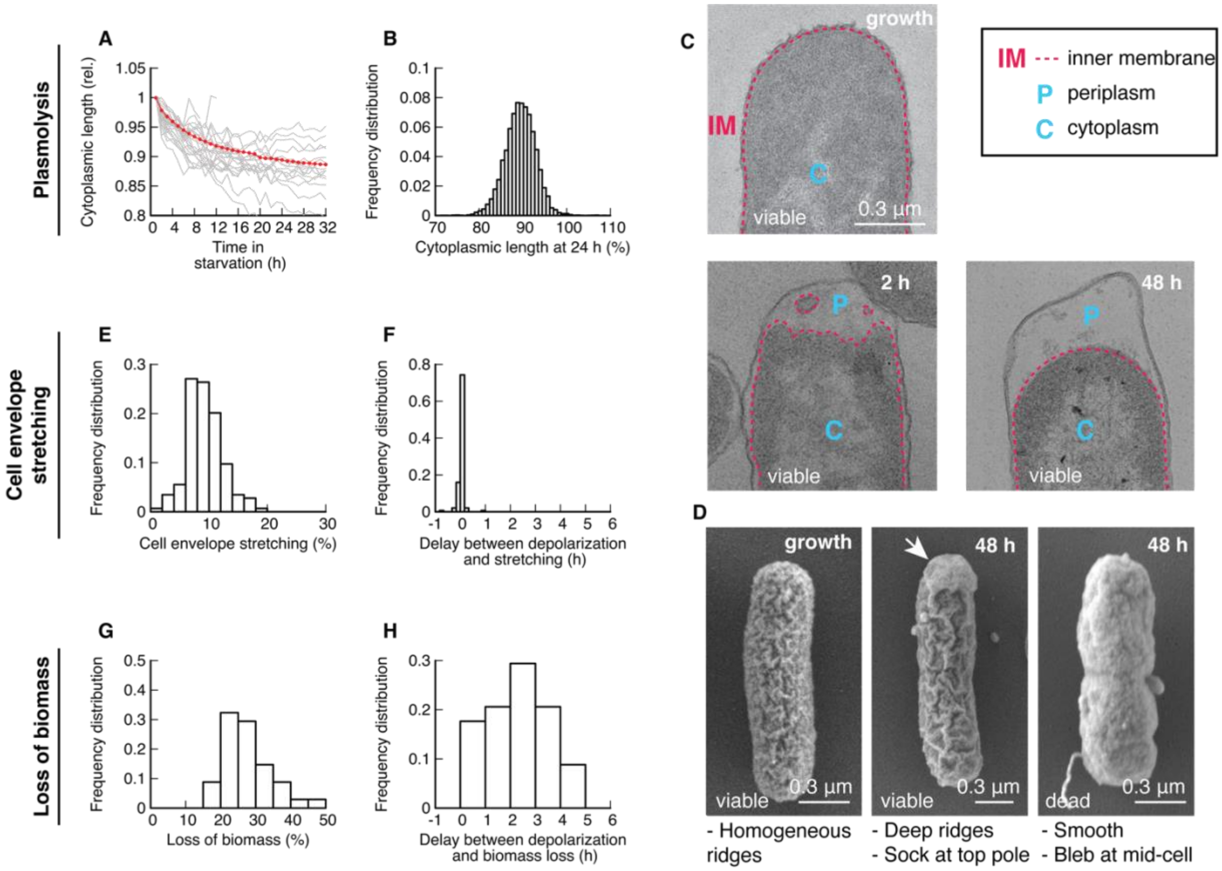


**Figure S 2 Characterization of cellular dynamics during starvation.** (**A**) Length of the cytoplasm of 25 individual bacteria measured by thresholding in phase contrast (grey, cut off 1 h before lysis) and the average of 14.498 bacteria (red). (**B**) Histogram of the contraction of the cytoplasm in individual bacteria at 24 h. Note that in the vast majority of bacteria (99.7%), the cytoplasm shrunk. (**C**) Transmission electron micrographs (TEM) of bacteria during exponential growth, 2 h and 48 h in starvation. The inner membrane (IM) is traced with a red dashed line. Periplasm (P), cytoplasm (C). Note that the inner membrane is closely aligned with the cell envelope during growth but retracts at 2 h in starvation. Excess material is visible in the undulating membrane and in blebs in the periplasm. At 48 h, excess membrane has disappeared. The near spherical shape of the inner membrane indicates that it is under tension. (**D**) Surface electron micrographs (SEM) of E. coli during growth show homogenous ridges of the cell envelope. In starvation the ridges on the surface are deeper, and a ‘sock-like’ structure is visible where the cytoplasm has contracted. In a dead cell, the surface is smooth, indicating that the surface structure is actively maintained. (**E**) Stretching of the cell envelope measured by thresholding in phase contrast. Cell envelope stretching is defined as the maximum length of the cell envelope divided by the average length of the cell envelope prior to expansion. (**F**) Delay between depolarization and stretching.

Swelling and depolarization coincide, $t_{\text{delay}} = (-0.05 \pm 0.5)$ h. (**G**) Loss of biomass, measured as the ratio of the average biomass up to 6 h prior to lysis and the average biomass 9 h after lysis. (**H**) Delay between loss of biomass (i.e., time-point of steepest slope of biomass data) and time-point of depolarization, $t_{\text{delay}} = (2.0 \pm 1.4)$ h.

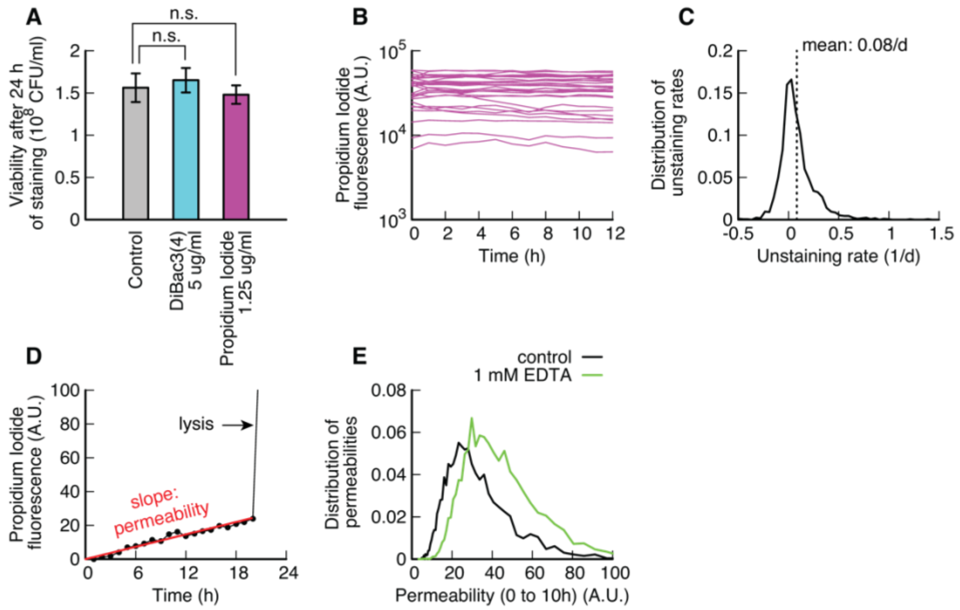


**Figure S 3 Propidium iodide and DiBac3(4) staining.** (**A**) Viability of E. coli after 24 h of starvation in the presence of either DiBac3(4) or Propidium Iodide at the indicated concentrations show no significant decrease in viability compared to an untreated control culture. (**B**) Propidium iodide staining dynamics of individual viable bacteria stained for 24 h in batch, washed and resuspended in fresh, carbon-free and stain-free medium. (**C**) Distribution of staining rates (slope of exponential fits to PI fluorescence signal) in carbon-free and stain-free medium. Mean unstaining rate is slow compared to the timescale of the experiment, meaning that staining can be considered irreversible. (**D**) Example of a PI staining time-trace. Lysis leads to a rapid increase of PI staining. Prior to lysis, we observe a slow increase of permeability. Because PI staining is irreversible, the slope of the absolute fluorescence signal is a measure for the total permeability of a bacterium. (**E**) Comparison of the permeability (measured between 0 and 10 h after entry to starvation) for a culture permeabilized with 1 mM EDTA compared to control.

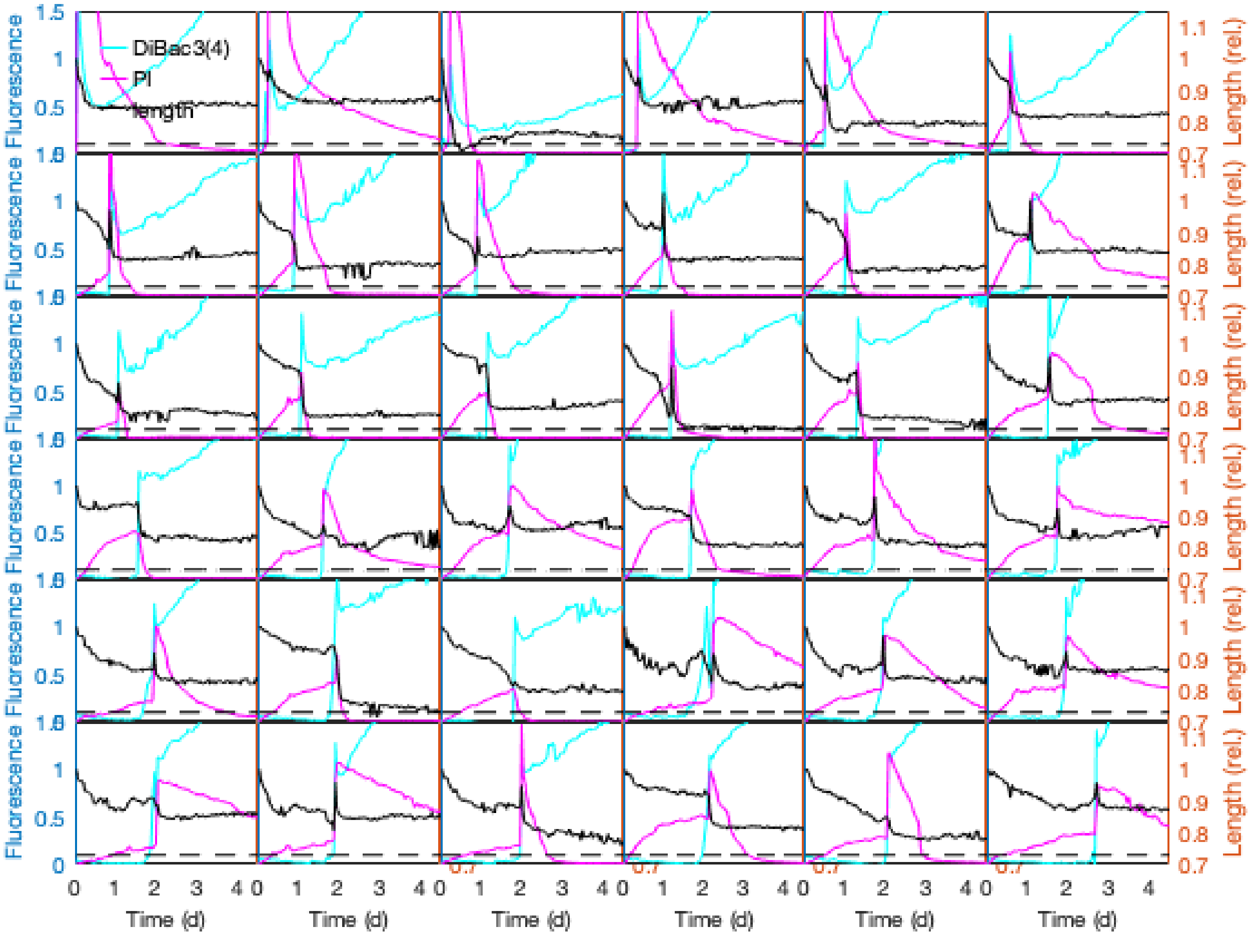


**Figure S 4 Single cell traces of depolarization and permeability in N+C- medium.** DiBac3(4) (depolarization, cyan) and PI (permeability, purple) and length (black, rel. to time-point zero. Right axis) for 36 randomly chosen bacteria. Note how the majority of bacteria show a sudden increase of DiBac3(4), length and PI, usually with only a short delay.

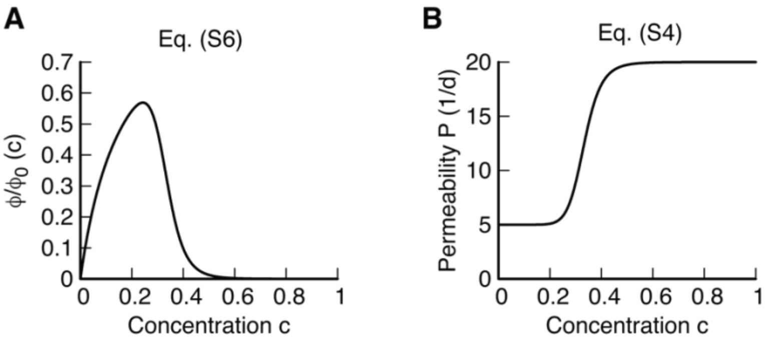


**Figure S 5 Model figure.** (**A**) Fraction of the maintenance rate that is used for pumping, see Eq. (S6). At low concentrations, pumping is limited due to the decrease in internal ion concentration, in a Michaelis-Menten-like function. At high concentration, pumping ceases because the membrane gets permeable, following a Hill equation. (**B**) In the model permeability increases from a base line to a higher value due to stretching of the membrane, see Eq. (S4).

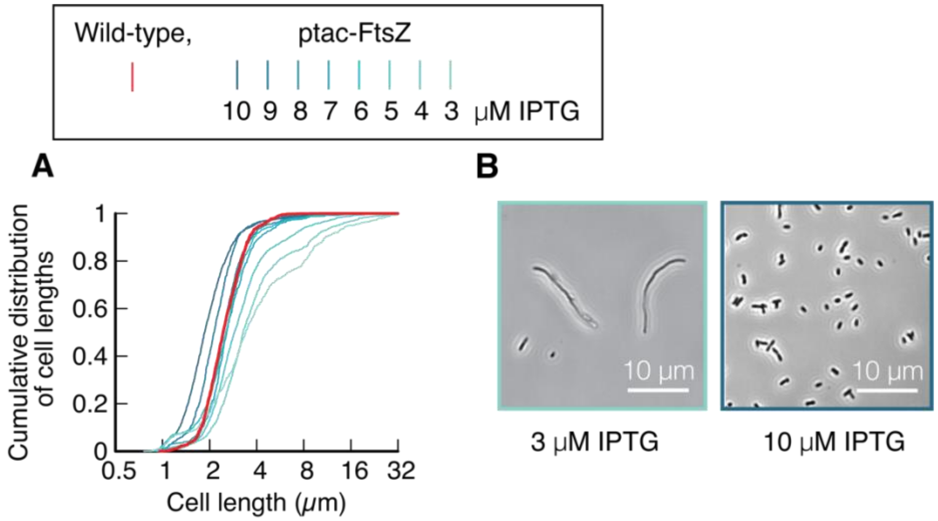


**Figure S 6 Titration of cell length with ptac-ftsZ, induced with IPTG during growth.** Dark shades of cyan indicate high expression of ftsZ, while bright shades indicate low expression of ftsZ. (**A**) Cumulative distribution of cell length for different induction levels (cyan) and wild-type (red). Note that at low induction levels the spread of the distribution increases. (**B**) Example images of the lowest and highest induction of FtsZ.

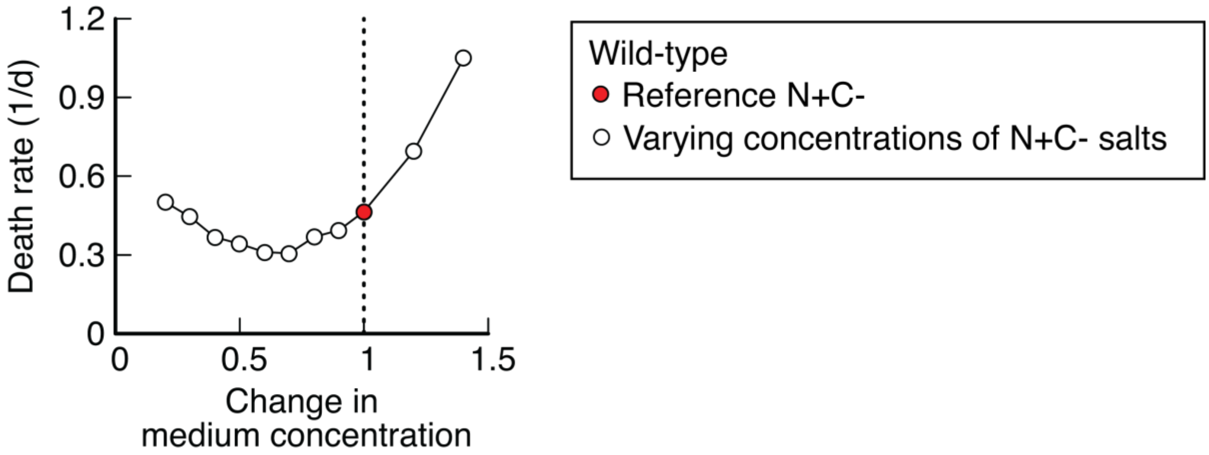


**Figure S 7 Effect of medium composition on death rate.** All cultures were grown in regular N+C+ medium supplemented with 10 mM glycerol and starved by switching medium. Reference: regular N+C- medium (red) n=3. Change in medium concentration means that the starvation medium was prepared with increased or decreased concentrations of all N+C- salts (white), n=1.

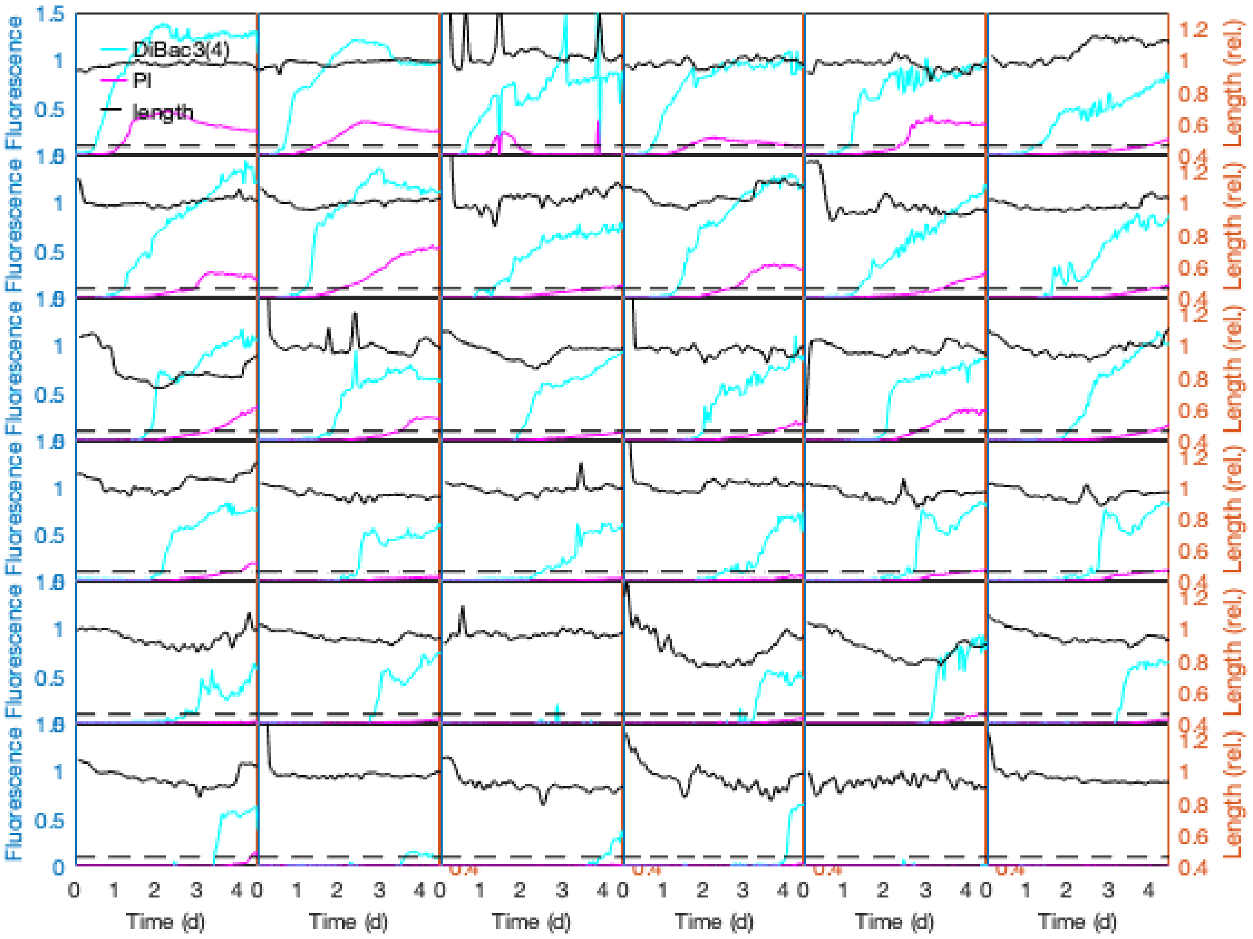


**Figure S 8 Single cell traces of depolarization, permeability and length in 'osmo-balanced' medium.** DiBac3(4) (depolarization, cyan), PI (permeability, purple) and length (Rel. to 20 to 30h, right y-axis) for 36 randomly chosen bacteria. Note how the majority of bacteria show a gradual increase of both DiBac3(4) and PI, usually with a delay of 1 to 2 days. Some bacteria divide shortly after entry to starvation, visible as a sharp drop in length. Because this data set has more noise, due to less well adherence of bacteria to the glass surface in this medium, we applied a Gaussian smoothing with 5 h window to the length measurement (black). This smoothing was not used in any of the fluorescence data and not used in the data presented in Fig. 4. Panels where no fluorescence traces are visible are bacteria that survived the displayed period.

### Supplementary Model

#### Introduction

In this section we describe the model that we used in the main text in Fig. 2. The model is based on the idea that the bacterium has to perform osmoregulation in order to remain viable. The goal of the model is to derive a) how the exponential decay of viability emerges b) what sets the time-scale and c) why individual bacteria are in plasmolysis, despite energy for pumping being limited.

In the model, we focus on osmoregulation, i.e., the pumping of ions across the cytoplasmic membrane that counteracts diffusive influx to prevent osmotic swelling and lysis.

#### Formulating the osmoregulation model

*Osmotic pressure*

Osmotic pressure is given by the concentration difference of all solutes between the inside and the outside of a membrane

$$\Pi = RT\left(\sum c_{\mathrm{in},i} - \sum c_{\mathrm{out},i}\right), \qquad \text{(S1)}$$

where $R$ is the gas constant and $T$ temperature. Many of the internal solutes are non-permeable, such as proteins, RNA, DNA and metabolites. These solutes, together with their positive counterions (because most macromolecules are net-negatively charged) form a set of solutes that are trapped in the cytoplasm of a viable bacterium, whose concentration we call $c_{\mathrm{biomass}}$. Solutes from the medium, such as $K^+$, $Na^+$ or $Cl^-$, on the other hand, can diffuse across the membrane. We do not know the precise nature of the permeability, and in particular we do not know the rate at which these metabolites permeate the membrane. But typically, any cell keeps the concentration of $Na^+$ low, while keeping $K^+$ high to balance charges from biomass. To avoid having to describe the dynamics of all ions, we instead subsume them into an effective ‘ion concentration’, that is outside $c_{\mathrm{out}}$ and inside $c_{\mathrm{in}}$, such that we can write osmotic pressure as

$$\Pi = RT(c_{\mathrm{biomass}} + c_{\mathrm{in}} - c_{\mathrm{out}}). \qquad \text{(S2)}$$

When the osmotic pressure in the cytoplasm becomes negative, e.g., if the bacterium is pumping ions out of the cytoplasm, the cytoplasm will contract such that the internal concentration with the new, reduced volume matches the outside concentration, $c_{\mathrm{biomass}} + c_{\mathrm{in}} = (n_{\mathrm{biomass}} + n_{\mathrm{in}})/V_{\mathrm{cytoplasm}} = c_{\mathrm{out}}$. This state of contracted cytoplasm is called plasmolysis, and the steady state maintenance of concentration gradients between inside and outside is called ion homeostasis.

As a simplification, we will not describe osmotic pressures and cytoplasmic volume changes explicitly in our model but instead focus only on the dynamics of the internal solutes. This removes the need to keep track of cytoplasmic volume changes. We further use an approximation of the internal concentration as $c_{\mathrm{in}} = n_{\mathrm{in}}/V_{\mathrm{cytoplasm,0}}$, where we normalize the internal ion abundance $n_{\mathrm{in}}$ by the volume of the cytoplasm in the non-contracted state, $V_{\mathrm{cytoplasm,0}}$.

*Ion homeostasis*

One fundamental problem of ion homeostasis is that ions can diffuse across the membrane and equilibrate the ion gradient. Thus, ion homeostasis requires transport of ions against the concentration gradient, which according to the second law of thermodynamics requires energy to be dissipated in the process. According to Fick’s law, the diffusive influx $j_{\mathrm{in}}$ is set by the permeability $P$ and ion gradient

$$j_{\mathrm{in}} = P(c_{\mathrm{out}} - c_{\mathrm{in}}). \quad \text{(S3)}$$

Here, we use the total influx of ions into over the bacterial surface, rather than using the definition of permeability being defined per area. Thus, permeability scales with cell surface area, $P \propto S$. For ion homeostasis to be maintained, this influx must be balanced with an outflux of equal magnitude, $j_{\mathrm{out}} = j_{\mathrm{in}}$.

Permeability of plasma membranes to inorganic ions increases if the membrane is stretched[1–3]. Because from TEM images we know that the cytoplasmic membrane in starvation is under tension (Fig. S2C) and we know any increase in internal solute concentration will lead to an increase of cytoplasmic volume and thus stretching of the membrane, we implemented stretch-activated permeability. Because stretching will increase with cytoplasmic volume, and cytoplasmic volume depends on the abundance of internal ions, $n_{\mathrm{in}}$, we expect permeability $P$ to be a function of $n_{\mathrm{in}}$ or alternatively $c_{\mathrm{in}} = n_{\mathrm{in}}/V_{\mathrm{cytoplasm,0}}$. Early in starvation, where there is an excess in membrane material (Fig. S2C), we see permeability stain PI increasing (Fig. S3D), so we suspect the functional form of $P$ to be constant for low $c_{\mathrm{in}}$ and increasing around some critical value when stretch-activation sets in, which is why we implemented a Hill function with constant offset,

$$P(c_{\mathrm{in}}) = P_0 + P_1 \frac{c_{\mathrm{in}}^n}{K_1^n + c_{\mathrm{in}}^n}. \tag{S4}$$

Permeability $P(c_{\mathrm{in}})$ for the parameters we picked in the model is shown in Fig. S5. Because in the simulation $c_{\mathrm{out}} = \mathrm{const}$, we wrote Eq. (S4) only as a function of $c_{\mathrm{in}}$.

From the experimental data it is clear that bacteria can regulate their permeability to some extent (Fig. 1F), that bacteria need to synthesize new lipids (Fig. 1G) and that bacteria consume lipids (Fig. 2C). While these are certainly interesting and important findings, we decided to not include them in the model to keep it simple and analytically tractable. Instead, we make the simplifying assumption that permeability for a given internal ion concentration is constant over time.

*Cannibalistic nutrient recycling*

To balance the diffusive influx of ions from the medium, the cell has to do active transport, which requires nutrients to be consumed. These nutrients can either come from a limited amount of storage or from recycling of biomass from perished bacteria. The latter, the ‘cannibalistic’ recycling, is a central aspect of starvation survival[4]. If we assume all bacteria to be equal, then we can describe the global nutrient resource $\nu$ as being supplied by the death of bacteria and consumed by viable bacteria. If $\alpha$ is the amount of nutrients, called the recycling yield[4], that can be recycled from a perished cell, then the nutrient supply will be $-\alpha\dot{N}$ where $-\dot{N}$ is the number of bacteria that die per time. Because bigger cells contain more nutrients, the recycling yield scales with cell volume, $\alpha \propto V$. Consumption on the other hand will be proportional to the number of viable bacteria $N$, and their maintenance rate $\beta$. Taken together, we can describe the dynamics of the nutrients as

$$\frac{\mathrm{d}\nu}{\mathrm{d}t} = -\alpha\frac{\mathrm{d}N}{\mathrm{d}t} - \beta N\ . \tag{S5}$$

If the nutrient in the culture is in steady state, i.e., $\dot{\nu} = 0$, then we can derive that maintenance rate is given by $\beta = \alpha\gamma$, where $\gamma = -\dot{N}/N$ is death rate.

We assume that a fraction $\phi$ of the maintenance rate is used for pumping ions against the gradient. To account for pumping to cease if $c_{\mathrm{in}}$ gets too small, we implement that $\phi(c_{\mathrm{in}})$ decreases linearly to zero at low concentrations following a Michaelis-Menten type function. At high concentrations, on the other hand, we suspect that once the membrane gets stretched, pumping ceases as ATP and other energy-rich molecules gradients disappear. To have this ceasing of pumping coincide with stretch-activated permeability, we choose the same dependence on $c_{\mathrm{in}}$

$$\phi(c_{\mathrm{in}}) = \phi_0\frac{c_{\mathrm{in}}}{K_2 + c_{\mathrm{in}}}\left(1 - \frac{c_{\mathrm{in}}^n}{K_1^n + c_{\mathrm{in}}^n}\right). \tag{S6}$$

$\phi(c_{\mathrm{in}})$ for the parameters we picked in the model is shown in Fig. S5.

We further absorb the proportionality constant of the cost of pumping (called $\kappa$ in the main text) into $\phi$ and write the active pumping flux of bacteria as

$$j_{\mathrm{pump}} = \alpha\phi(c_{\mathrm{in}})\gamma. \tag{S7}$$

*Dynamics of cytoplasmic solutes*

Using the combination of diffusive influx and active pumping, we can calculate the dynamics of the internal ion concentration $c_{\mathrm{in}}$.

$$\frac{\mathrm{d}c_{\mathrm{in}}}{\mathrm{d}t} = P(c_{\mathrm{in}})(c_{\mathrm{out}} - c_{\mathrm{in}}) - j_{\mathrm{pump}} + \xi(t) \tag{S8}$$

In addition, we assume that there is noise in the underlying biological processes that affect permeability or pumping, which we subsume in a normally distributed Wiener process with diffusion constant $D$. Plugging in Eq. (S7) we get

$$\frac{\mathrm{d}c_{\mathrm{in}}}{\mathrm{d}t} = P(c_{\mathrm{in}})(c_{\mathrm{out}} - c_{\mathrm{in}}) - \alpha\phi(c_{\mathrm{in}})\gamma + \xi(t). \quad \text{(S9)}$$

*Death*

Finally, to conclude Eq. (S9), we need to define when a bacterium is called dead, in order to calculate death rate. Because death is due to lysis (Fig. 1), which is in turn due to an increase in internal ion concentration, we call a bacterium dead if there is a significant increase of the internal ion concentration, $c_{\mathrm{in}}/c_{\mathrm{out}} \gg 0$ that goes beyond just stretching of the cytoplasmic membrane $c_{\mathrm{in}} > K_1$.

*Pseudo-potential*

One way of conceptualizing the dynamics of the system is mapping the problem onto a potential $U(c_{\mathrm{in}})$, where

$$\frac{\mathrm{d}c_{\mathrm{in}}}{\mathrm{d}t} = -U'(c_{\mathrm{in}}) + \xi(t). \quad \text{(S10)}$$

In a physics setting, $-U'(c_{\mathrm{in}})$ would be the restoring force acting on $c_{\mathrm{in}}$. As we will calculate below, the potential can have a local minimum at low $c_{\mathrm{in}}$, where noise $\xi(t)$ can push the cell out and across the barrier towards high $c_{\mathrm{in}}$ and thus lysis.

To understand the dynamics, we will discuss the shape of $U$. By looking at the zeros of $U'(c_{\mathrm{in}})$ we can understand the maxima and minima of the system. For small $c_{\mathrm{in}}$,

$$U'(c_{\mathrm{in}}) \approx -Pc_{\mathrm{out}} + \frac{\alpha\phi_0 c_{\mathrm{in}}\gamma}{K_2} \quad \text{for } c_{\mathrm{in}} \ll c_{\mathrm{out}}, \quad \text{(S11)}$$

i.e., negative for $c_{\mathrm{in}} = 0$, but increasing. $U'(c_{\mathrm{in}})$ can become positive if the linear term outweighs the constant term before pumping is shut-off by stretch-activated permeability around $c_{\mathrm{in}} < K_1$; see Eq. (S6). This the case for $Pc_{\mathrm{out}}K_2/\alpha\phi_0\gamma > K_1$, which depends on death rate $\gamma$.

In the regime of $c_{\mathrm{in}} \approx c_{\mathrm{out}}$, $\phi(c_{\mathrm{in}}) = 0$ and $P = P_0 + P_1$. Thus

$$U'(c_{\mathrm{in}}) \approx -(P_0 + P_1)(c_{\mathrm{out}} - \mathrm{c_{in}}) = 0 \quad \text{for } c_{\mathrm{in}} = c_{\mathrm{out}} \tag{S12}$$

At this zero, the second derivative $U''(c_{\mathrm{in}}) = (P_0 + P_1) > 0$, meaning that it is always a minimum of $U(c_{\mathrm{in}})$.

Thus, the potential $U(c_{\mathrm{in}})$ will always have one zero at $c_{\mathrm{out}} = \mathrm{c_{in}}$, and can have in addition one minimum at a low concentration and a maximum at an intermediate concentration.

*Kramers' escape from the potential well*

If the potential has two minima, then the noise-mediated escape from the minimum at concentration $\mathrm{c_{min}}$ across the barrier at $c_{\mathrm{max}}$ can be understood with Kramers' escape theory[5,6]. If the concentration is in a quasi-steady state at low $c_{\mathrm{in}}$, and the barrier $E_{\mathrm{b}} = U(c_{\mathrm{max}}) - U(\mathrm{c_{min}})$ is bigger than the diffusion constant $E_{\mathrm{b}} > D$, then the escape rate can be approximated as

$$k \approx \frac{1}{2\pi}[U''(\mathrm{c_{min}})|U''(c_{\mathrm{max}})|]^{\frac{1}{2}} \exp\left(\frac{E_{\mathrm{b}}}{D}\right) \tag{S13}$$

In our model we equate the escape rate and the death rate, $k = \gamma$, meaning that any cell that crosses the barrier has irreversibly died. But because the potential itself depends on the death rate, see Eq. (S9) and (S10), this leads to the interesting system where the escape rate across the barrier determines the height of the barrier.

*Emergence of a steady-state death rate and exponential decay*

The feedback loop in which the escape rate across the barrier affects the barrier height brings about a self-adjusting system with a unique steady-state death rate. We restrict ourselves here to cases where, in addition to the requirements for Kramers escape theory, we can assume that the concentrations at which the pseudo-potential achieves its minimum and maximum do not significantly change with death rate and can be approximated as constants. While this assumption breaks down with very large fluctuations in the death rate, it is reasonable given the small fluctuations that usually occur.

We define the function

$$f(c_{\text{in}}) \equiv P(c_{\text{in}})(c_{\text{out}} - c_{\text{in}}) - j_{\text{pump}} = -U'(c_{\text{in}}). \quad \text{(S14)}$$

This means that Eq. (S13) is equivalent to

$$k \approx \frac{1}{2\pi}[-f'(\mathrm{c}_{\min})f'(c_{\max})]^{\frac{1}{2}} \exp\left(\frac{\int_{c_{\min}}^{c_{\max}} f(c)\mathrm{d}c}{D}\right) \quad \text{(S15)}$$

Noting that form of $j_{\text{pump}}$ in Eq. (S7) depends linearly on the death rate, we can define functions $f_{\text{influx}}$ and $f_{\text{pump}}$ independent of the death rate such that

$$\mathrm{f}(c_{\text{in}}) = \mathrm{f}_{\text{influx}}(c_{\text{in}}) - f_{\text{pump}}(c_{\text{in}})\gamma. \quad \text{(S16)}$$

Then, if we define the constants

$$\begin{aligned} A_0 &\equiv -f'_{\text{influx}}(c_{\min}), \\ A_1 &\equiv f'_{\text{pump}}(c_{\min}), \\ B_0 &\equiv f'_{\text{influx}}(c_{\max}), \\ B_1 &\equiv -f'_{\text{pump}}(c_{\max}), \\ C_0 &\equiv \tfrac{1}{\mathrm{D}}\textstyle\int_{c_{\min}}^{c_{\max}} f_{\text{influx}}(c)\mathrm{d}c, \text{ and} \\ C_1 &\equiv \tfrac{1}{D}\textstyle\int_{c_{\min}}^{c_{\max}} f_{\text{pump}}(c)\mathrm{d}c, \end{aligned} \quad \text{(S17)}$$

we can rewrite the Kramers escape rate as

$$\mathrm{k} \approx \tfrac{1}{2\pi}[(A_0 + A_1\gamma)(B_0 + B_1\gamma)]^{\frac{1}{2}} \exp(\mathrm{C}_0 - \mathrm{C}_1\gamma). \quad \text{(S18)}$$

The physical constraints of the system can then impose qualitative restrictions on the behavior of the escape rate as a function of the death rate. Because we are working in a regime where $c_{\min}$, the equilibrium corresponding to healthy cells, is low enough that there is very little stretch-induced permeability while pumping is still increasing, we can conclude that $A_0$ is positive since the permeability will remain relatively constant while the concentration gradient will decrease if the internal ion concentration increases, and $A_1$ is positive since the pumping rate will increase with greater ion concentration. Similarly, since the maximum occurs where stretch-induced permeability is very high and pumping becomes more difficult, $B_0$ and $B_1$ are both positive. Finally, since influx and pumping are always positive whenever $c_{\text{in}}$ is positive, $C_0$ and $C_1$ must both be positive. Under these conditions, the graph of $k$ as a function of γ is always defined, infinitely differentiable, and positive for nonnegative γ, and it approaches zero as γ approaches infinity.

Steady-state death rates are found by finding values such that

$$k(\gamma) = \gamma. \tag{S19}$$

Using the expression for $k$ found in Eq. (S18) and rearranging, we find that this requirement is equivalent to

$$2\pi\gamma[(A_0 + A_1\gamma)(B_0 + B_1\gamma)]^{-1/2} = \exp(C_0 - C_1\gamma). \tag{S20}$$

Because

$$\frac{\mathrm{d}}{\mathrm{d}\gamma}\left[2\pi\gamma[(A_0 + A_1\gamma)(B_0 + B_1\gamma)]^{-1/2}\right] = \frac{\pi(2A_0B_0 + A_0B_1\gamma + A_1B_0\gamma)}{[(A_0 + A_1\gamma)(B_0 + B_1\gamma)]^{3/2}}, \tag{S21}$$

which is always positive for nonnegative γ, Eq. (S20) requires that an increasing function of γ must intersect with a decreasing function to achieve equilibrium. Furthermore, since the LHS is less than the RHS for very small γ and greater than the RHS when γ is large, there exists a unique death rate $\gamma^*$ at which these two curves intersect.

Under perturbations from the steady-state death rate $\gamma^*$, the system will adjust itself back toward equilibrium. The above analysis implies that $k > \gamma$ if $\gamma < \gamma^*$ and $k < \gamma$ if $\gamma > \gamma^*$, which means that the rate of escape from the pseudo-potential will correct itself toward $\gamma^*$ if the death rate at any moment is above or below the steady-state death rate. Thus, following random fluctuations or changes to the death rate by outside factors, the system will relax back into the steady-state equilibrium.

*Stochastic implementation of the model*

To run a stochastic simulation of the model, we implemented noise as a normal-distributed contribution to concentration of every cell $i$, where at each time-step $c_t^i = c_{t-1}^i + \Delta t \frac{\mathrm{d}c^i}{\mathrm{d}t} +$

$\xi(D, \Delta t)\Delta$, where $\xi(D, \Delta t)$ is a random, normal distributed number with mean 0 and standard deviation $\sqrt{2D\Delta t}$. In addition, we simulated the nutrient resource for each individual cell by consuming all available nutrients at each time-step $\nu_{t-1} = \beta\Delta t$, and adding nutrients of the number of bacteria that have died in the time-step, $-\Delta N$, proportionally to the remaining fraction of viable bacteria, $\nu_t = -\alpha\, \Delta N/N$.

*Parameters of the model*

In the simulation we used the following parameters:

| **Parameter** | **Comment** | **Value** | **Unit** |
|---|---|---|---|
| $c_{\mathrm{out}}$ | All concentrations normalized to $c_{\mathrm{out}}$ | 1 | 1 |
| $P_0$ | Permeability times surface area, unstretched | 5 | $\mathrm{d}^{-1}$ |
| $P_1$ | Permeability times surface area, stretched-activated contribution | 15 | $\mathrm{d}^{-1}$ |
| $n$ | Hill coefficient in Eq. (S4) and (S6) | 10 | - |
| $K_1$ | Half-maximum of stretch-activation | 1/3 | $c_{\mathrm{out}}$ |
| $K_2$ | Half-maximum of pumping | 1/6 | $c_{\mathrm{out}}$ |
| $\alpha\phi_0$ | Recycled nutrients that are used for pumping, divided by cost of pumping | 16 | $c_{\mathrm{out}}$ |
| $D$ | Diffusion coefficient | 0.1 | $\frac{c_{\mathrm{out}}^2}{\mathrm{d}}$ |
| $c_{\mathrm{tresh}}$ | Threshold at which a cell is called dead and its nutrients are distributed. | 0.35 | $c_{\mathrm{out}}$ |
